# IMGT® at scale: FAIR, Dynamic and Automated Tools for Immune Locus Analysis

**DOI:** 10.1101/2025.08.12.669823

**Authors:** Gaoussou Sanou, Guilhem Zeitoun, Taciana Manso, Milad Eidi, François Grand, Anjana Kushwaha, Myriam Croze, Chahrazed Debbagh, Axel Vaillant, Maria Georga, Ariadni Papadaki, Ifigeneia Sideri, Shamsa Batool, Turkan Samadova, Joumana Jabado-Michaloud, Géraldine Folch, Véronique Giudicelli, Patrice Duroux, Sofia Kossida

## Abstract

IMGT®, the international ImMunoGeneTics information system®, has advanced its comprehensive platform for the analysis of immunoglobulin (IG) and T cell receptor (TR) genes through the development of new automated and scalable tools. This article presents major updates aligned with IMGT’s three axes of research. Axis I introduces dynamic resources such as IMGT/GeneTables, IMGT/AssemblyComparison, and IMGT/StatAssembly, enabling real-time access to annotated genomic data and quality assessment of assemblies. Axis II enhances repertoire analysis with a redesigned IMGT/GeneFrequency tool and new customization features in IMGT/V-QUEST, supporting flexible exploration of IG and TR gene expression. Axis III improves the accurate prediction of peptide-MHC thanks to IMGT/RobustpMHC. Additionally, the IMGT Knowledge Graph (IMGT-KG) and its therapeutic extension, IMGT/mAb-KG, provide semantically structured access to more than 100 million immunogenetic triplets, integrating IMGT databases and linking IMGT content to external biomedical resources. These developments promote standardization, interoperability, and integrative analysis across immunogenetics and clinical applications, reinforcing IMGT’s role as a core reference in the era of FAIR data and personalized medicine.

## 1 Introduction

IMGT®, the international ImMunoGeneTics information system® offers a standardized and comprehensive platform for analysing and understanding immunoglobulins (IG, antibodies) and T cell receptors (TRs) [1]. IMGT integrates high-quality curated databases [2], advanced bioinformatics tools, and a unified vocabulary to ensure consistent and accurate immune repertoire analysis [3]. IMGT includes several specialized databases [4], such as IMGT/LIGM-DB for nucleotide sequences, IMGT/GENE-DB for genes and alleles [5], IMGT/2Dstructure-DB and IMGT/3Dstructure-DB for amino acid sequences and their 2D and 3D structures [6], and IMGT/mAb-DB dedicated to therapeutic monoclonal antibodies and their clinical applications [7]. To address the evolving needs of the community and overcome specific challenges in immunogenetics research, IMGT has developed new tools and resources with specialized capabilities for analysing, visualizing, and interpreting adaptive immune system data. The development of these resources and tools is driven by three key axes of research and development [8]. Each axis introduces new resources or tools designed to address specific research challenges, thereby expanding the overall functionality of the IMGT workflow.

Axis I of IMGT is dedicated to deciphering the adaptive immune response by analysing the organization and structure of immunoglobulin (IG) and T cell receptor (TR) loci across jawed vertebrates. This fundamental research involves identifying and characterizing IG and TR genes and alleles, which are crucial for understanding the genetic basis of immune diversity. A significant achievement of Axis I is the extraction of complete IG and TR loci from genome assemblies, facilitating a comprehensive understanding of the immune repertoire. The resulting data are used to update IMGT databases such as IMGT/LIGM-DB and IMGT/GENE-DB. Within this axis, data can be visualized using various web resources that compile annotated data and provide knowledge pages, such as those found in the IMGT Web Resources: IMGT Repertoire. Historically, these webpages were static and manually generated. Recently, several have been automated and new dynamic webpages have been created, including IMGT/GeneTable, IMGT/MultipleGenomeViewer, IMGT/ComparisonAssembly, IMGT/StatAssembly, and IMGT/CDRLengths.

The data generated through Axis I are compiled into IMGT reference directories, which serve as essential resources for the subsequent axes of research and development. These directories provide standardized references that support the exploration of expressed IG and TR repertoires in Axis II and analysis of the structural aspects of adaptive immune proteins in Axis III.

Axis II focuses on investigating IG and TR gene diversity and expression patterns, which are essential for understanding adaptive immune responses in both physiological and pathological contexts. It provides indispensable tools for the analysis of expressed IG and TR sequences, enabling studies on V(D)J recombinations, somatic hypermutations, and clonal expansions based on nucleotide sequences using IMGT/V-QUEST and its high throughput version IMGT/HighV-QUEST. Additionally, further analytical capabilities are offered by IMGT/StatClonotype and IMGT/JunctionAnalysis. IMGT/GeneFrequency has been enhanced and provides new features and functionalities for an intuitive way to explore the gene frequencies in different species. In addition, IMGT/V-QUEST has been enhanced with new features allowing the customization of the reference directory set.

The last research axis of IMGT, Axis III, focuses on analysing the two-dimensional (2D) and three-dimensional (3D) structures of adaptive immune proteins, including IG and TR. By emphasizing the structural analysis of these proteins, Axis III enhances our understanding of the molecular mechanisms underlying immune responses. In addition to the existing tools in this axis, such as IMGT/DomainGapAlign and IMGT/Collier-de-Perles, IMGT has developed IMGT/RobustpMHC [9], a new machine learning-based tool to predict peptide-MHC (major histocompatibility complex) interactions, a crucial process for immune recognition.

The design and content of the IMGT databases, along with the tools and web resources, are the result of an extensive process of curation and standardization of immunogenetics knowledge, facilitated by IMGT-ONTOLOGY [3]. To bridge the gap between nucleotide and amino acid sequence databases, IMGT created the IMGT Knowledge Graph (IMGT-KG) [10], a comprehensive resource integrating the five IMGT databases and connecting to external related biomedical resources including Open Biological and Biomedical Ontology (OBO) Foundry resources or National Cancer Institute Thesaurus (NCIt) [11]. Building on this foundation, IMGT/mAb-KG, is the IMGT-KG for therapeutic monoclonal antibodies (mAb) providing access to knowledge about the mAbs, their structures, their targets, the related clinical indications and their mechanism of action [12].

In this article, we highlight the latest advancements in IMGT, developed through its three research and development axes, which collectively establish IMGT® as an indispensable system for studying the adaptive immune system, assisting in the design of therapeutic interventions, and paving the way for personalized medicine.

## 2 Axis 1 : Dynamic Web Resources

### 2.1 IMGT/GeneTables

IMGT/GeneTables provides access to the repertoire of genes and alleles for a given species and gene type within a locus (IGHV, IGHD, etc). For that, the IMGT/GeneTables separates the IMGT reference sequences from IMGT literature sequences [5]. The IMGT/GeneTables provides information about the alleles, including the functionality, the clone name, the rearranged or transcribed information, the accession number, and the sequence associated to the alleles. Additionally, bibliographic references and specific notes regarding alleles are included in this table. IMGT/GeneTables is part of [IMGT Repertoire](https://imgt.org/IMGTrepertoire/), the global immunogenetics web resources providing access to expert-curated data on the IG, TR, major histocompatibility (MH) and related proteins of the immune system (RPI). [8]. Until recently the gene table for each species and gene type were generated manually following locus annotation. This approach was time-consuming, susceptible to human error, and may quickly become outdated when new annotations or modifications were introduced.

Recently, the [IMGT/GeneTables](https://www.imgt.org/IMGTrepertoire/LocusGenes/genetable/autotable.php) has been automated and data from more than 100 gene types across more than 30 species are now available to the public. The data are extracted in real-time from IMGT databases, ensuring consistency and reducing errors. The design has been updated to include new information, such as the IMGT allele confidence score, which reflects the number of literature-reported sequences available for a given allele. The data can be downloaded as an Excel file. In species such as mouse, it is also possible to identify alleles that are specific to a single strain. Figure 1 provides a visualization example of the locus IGHJ in *Homo sapiens*.

**Figure 1:**
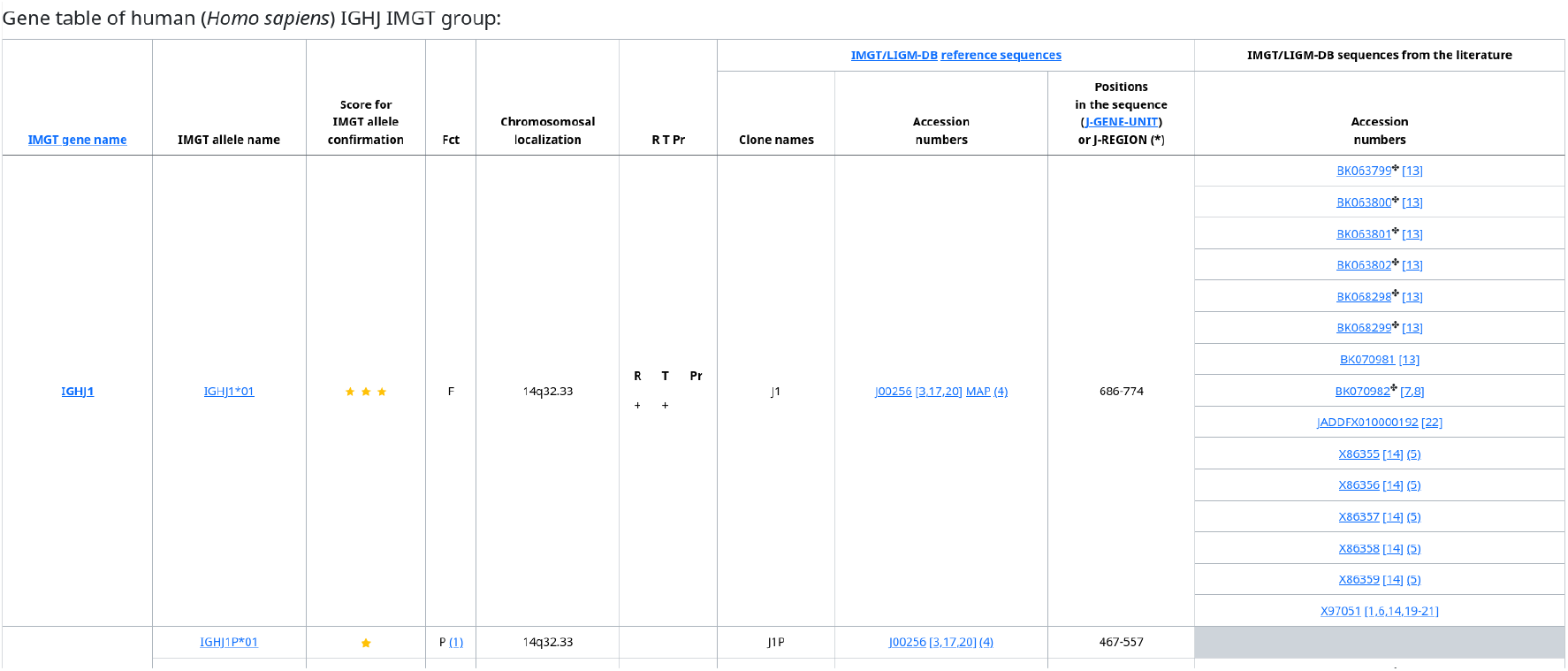
Visualisation of IGHJ locus of *Homo sapiens* as of 22 May 2025, accessible from [IMGT webpage](https://imgt.org/IMGTrepertoire/LocusGenes/genetable/autotable.php?species=Homo+sapiens&group=IGHJ).

### 2.2 IMGT/AssemblyComparison

IMGT offers a wide range of annotated assemblies across different species. As of 17 July 2025, there are for instance, 12 assemblies annotated for human, 11 for dogs and seven for mouse. These different assemblies can reveal gene variations, such as copy number variations (CNVs) [13] or differences in gene functionalities. To facilitate their comparison, we developed [IMGT/AssemblyComparison](https://www.imgt.org/IMGTrepertoire/LocusGenes/locusdesc/assembly_compare.php), a new dynamic web interface for exploring locus gene repertoires. IMGT/AssemblyComparison is an automated tool that, for a given species and locus, displays all genes by subgroup and functionality, based on IMGT-annotated and localized assemblies from [IMGT/GENE-DB genomic localizations](https://www.imgt.org/genedb/#resultPage_localizations). The tool identifies shared and distinct genes, and variations in allele functionality. Data are retrieved in real time from IMGT databases to generate the assembly comparison. In the output (Figure 2), functional differences between alleles of the same gene are highlighted, along with shared genes and other detected variations. The percentage of genes per functionality is also displayed as a bar chart. Additional information is provided in a pop-up window and a summary table at the bottom of the page, which can be downloaded as a CSV file (Figure 2).

**Figure 2:**
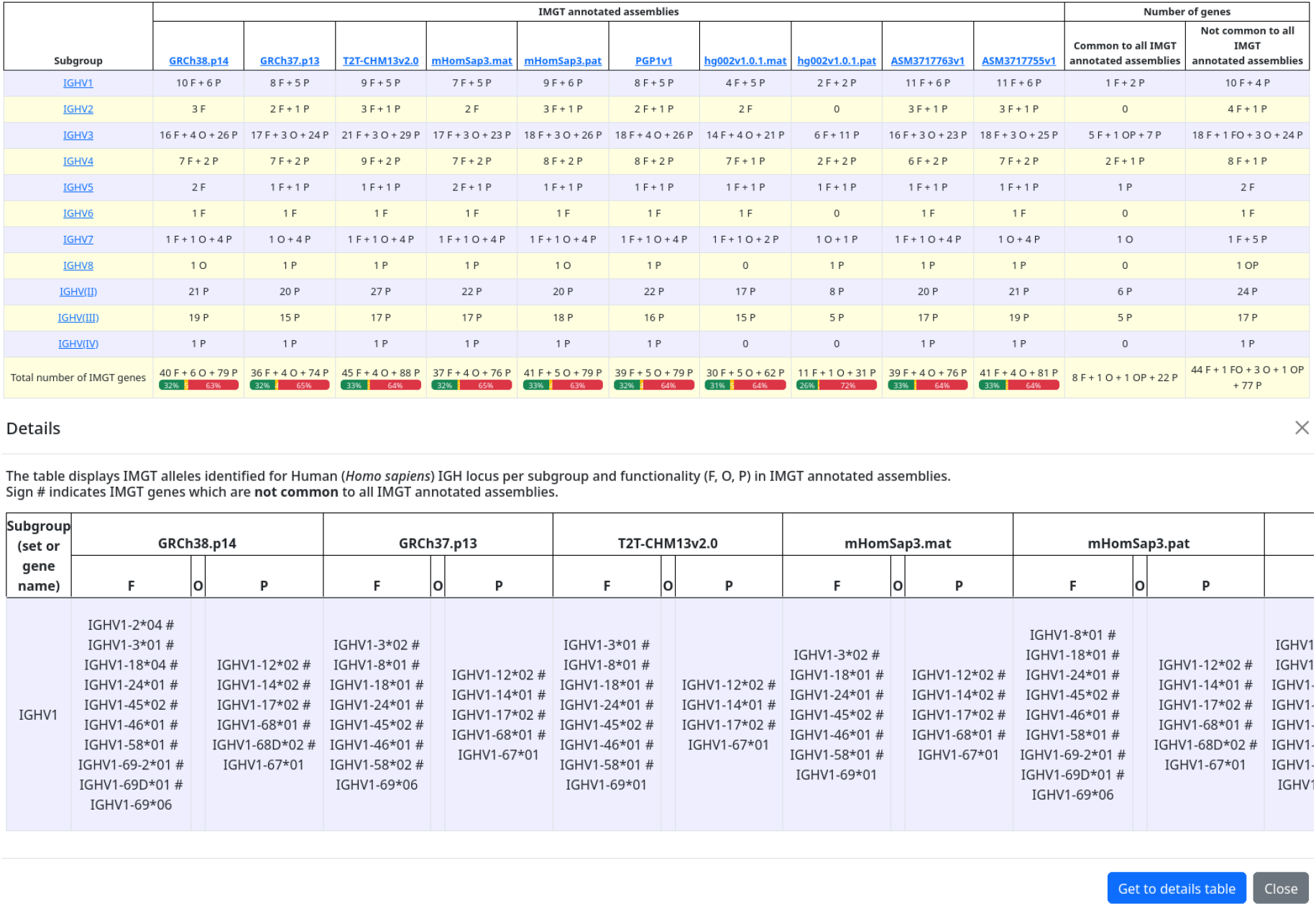
Visualization of the IGH locus assembly in *Homo sapiens* as of 6 August 2025, accessible via the [IMGT website](https://imgt.org/IMGTrepertoire/LocusGenes/locusdesc/assembly_compare.php?species=Homo+sapiens&locus=IGH) (top). Details are shown for the IGHV1 subgroup of Homo sapiens, including one functional gene (IGHV1-69) and two pseudogenes (IGHV1-67 and IGHV1-68), all conserved across IMGT-annotated and localized assemblies.

### 2.3 IMGT/CDRLengths

IG and TR variable genes are categorized in subgroups [14] based on the similarity of their coding region (V-REGION). The length of the complementarity-determining regions (CDR) of the variable genes is an important factor contributing to gene diversity. In recognition of this, we introduce *[IMGT/CDRLengths](https://www.imgt.org/IMGTrepertoire/2D-3Dstruct/FClength/cdr_home.php)*, dynamic web pages that visualise the diversity of CDR lengths within each subgroup [1] as well as differences among alleles. The data are also retrieved in real time from IMGT databases and can be downloaded as a CSV file (Figure 3).

**Figure 3:**
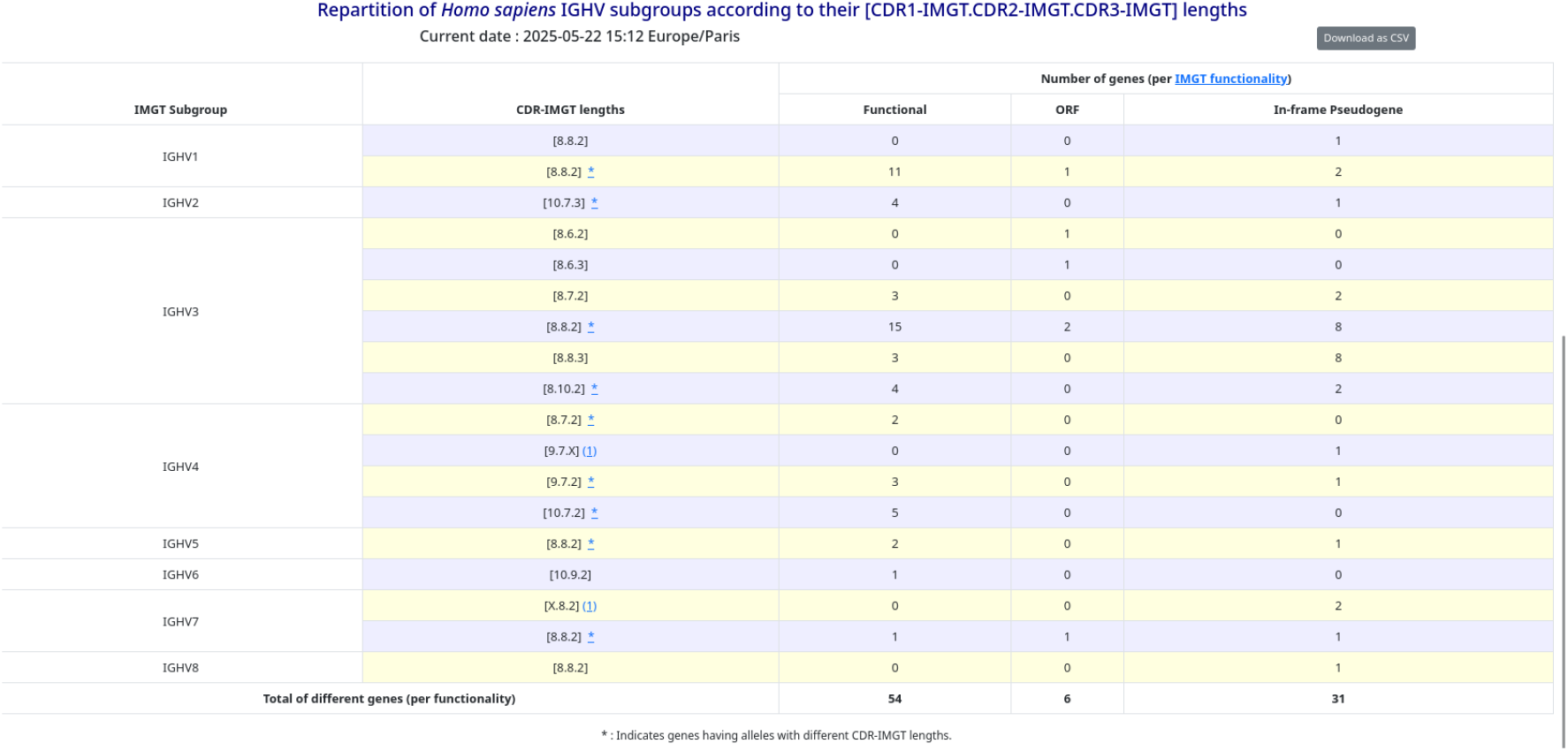
Visualisation of CDR lengths for the IGH locus of *Homo sapiens* as of 22 May 2025, accessible from [IMGT webpage](https://www.imgt.org/IMGTrepertoire/2D-3Dstruct/FClength/cdr_home.php?species=Homo+sapiens&locus=IGH).

### 2.4 IMGT/MultipleGenomeViewer

Inspired by the Richardson et al. work [15], we implemented a new version of the Multiple Genome Viewer (MGV), IMGT/MultipleGenomeViewer (IMGT/MGV), based on IMGT/GENE-DB genomic localizations. In addition to displaying all IG/TR genes and loci on the chromosome for each chosen assembly, IMGT/MGV facilitates position retrieval, gene and allele comparison, functionality overview, and the comparison of assemblies within the same species (Figure 4). Sequences can be downloaded in a FASTA file. As of March 5th, 2025, data for four species are available.

**Figure 4:**
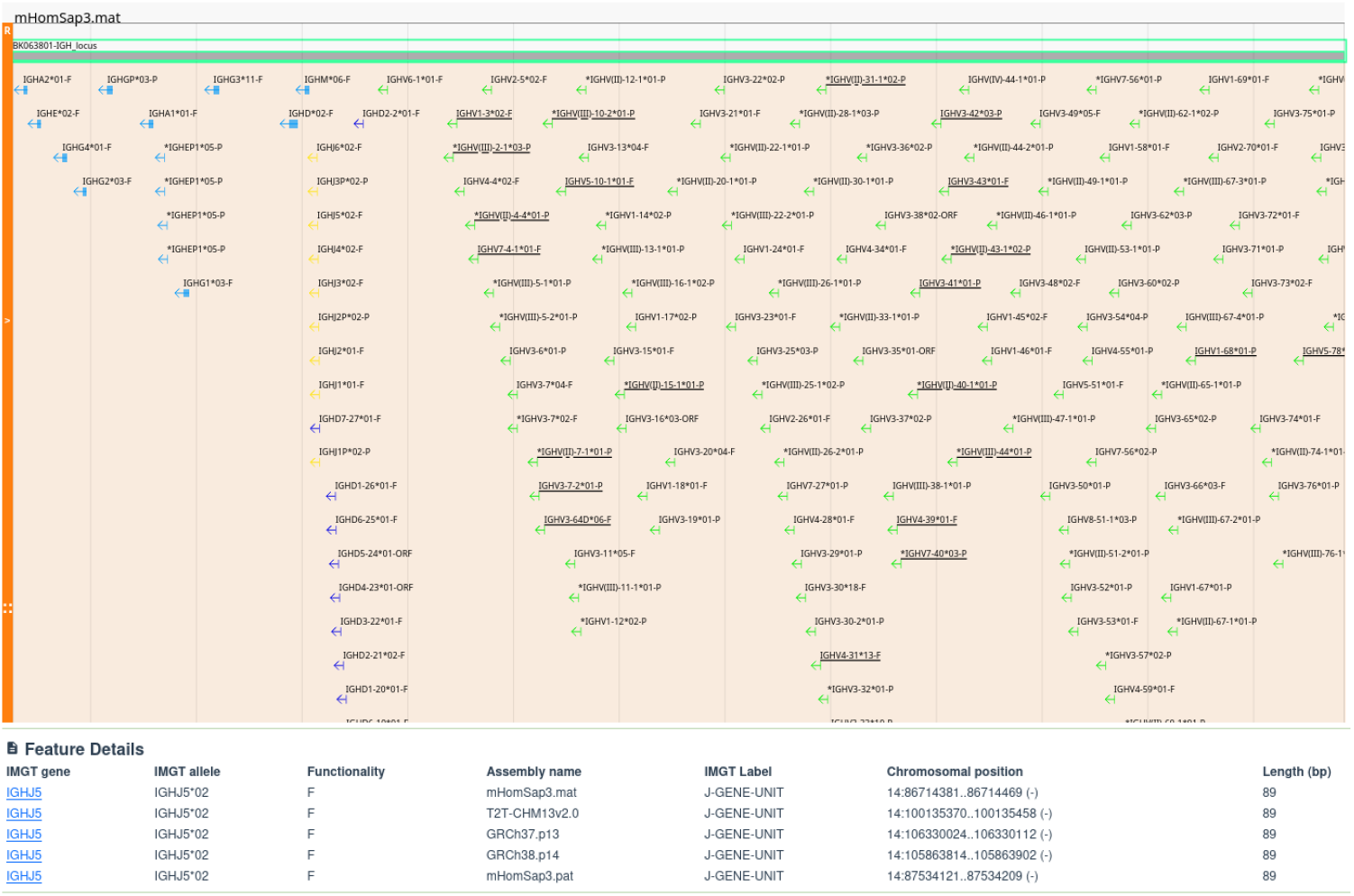
Visualisation of the mHomSap3.mat (*Homo sapiens*) assembly on IMGT/MGV as of 5 March 2025 (top). The lower panel shows the list of IGHJ5 alleles across human assemblies, along with their position within each locus.

### 2.5 IMGT/StatAssembly

#### 2.5.1 Description of the tool

Inspired by the work of Zhu et al. on the CloseRead tool [16], IMGT/StatAssembly [17] allows the analysis of an alignment file in BAM format. This file can be generated using the minimap2 tool [18] and should contain either a CS, MD or CIGAR tag =/X (match/mismatch distinction).

IMGT/StatAssembly takes as input the positions of the IG and/or TR loci, either extracted from the IMGT Locus Description (IMGT Repertoire) or identified through sequence similarity. The tool then analyses each IG/TR locus region provided and counts the number of reads based on their mapping score (Figure 5). Regions covered by fewer than three primary reads are labeled as ‘break’ positions. Secondary^1^ and supplementary^2^ alignments are displayed, along with the number of overlapping primary reads (i.e., reads covering the given position and its adjacent positions (Figure 5). The number of overlapping primary reads should be close to, if not equal to, the total number of primary reads. A decrease may indicate regions where reads fail to overlap, which could signal potential assembly issues. High-quality assemblies are expected to show consistent coverage with a predominance of green (high mapping quality) reads and minimal numbers of secondary, supplementary, or low-mapping-quality primary reads.

**Figure 5:**
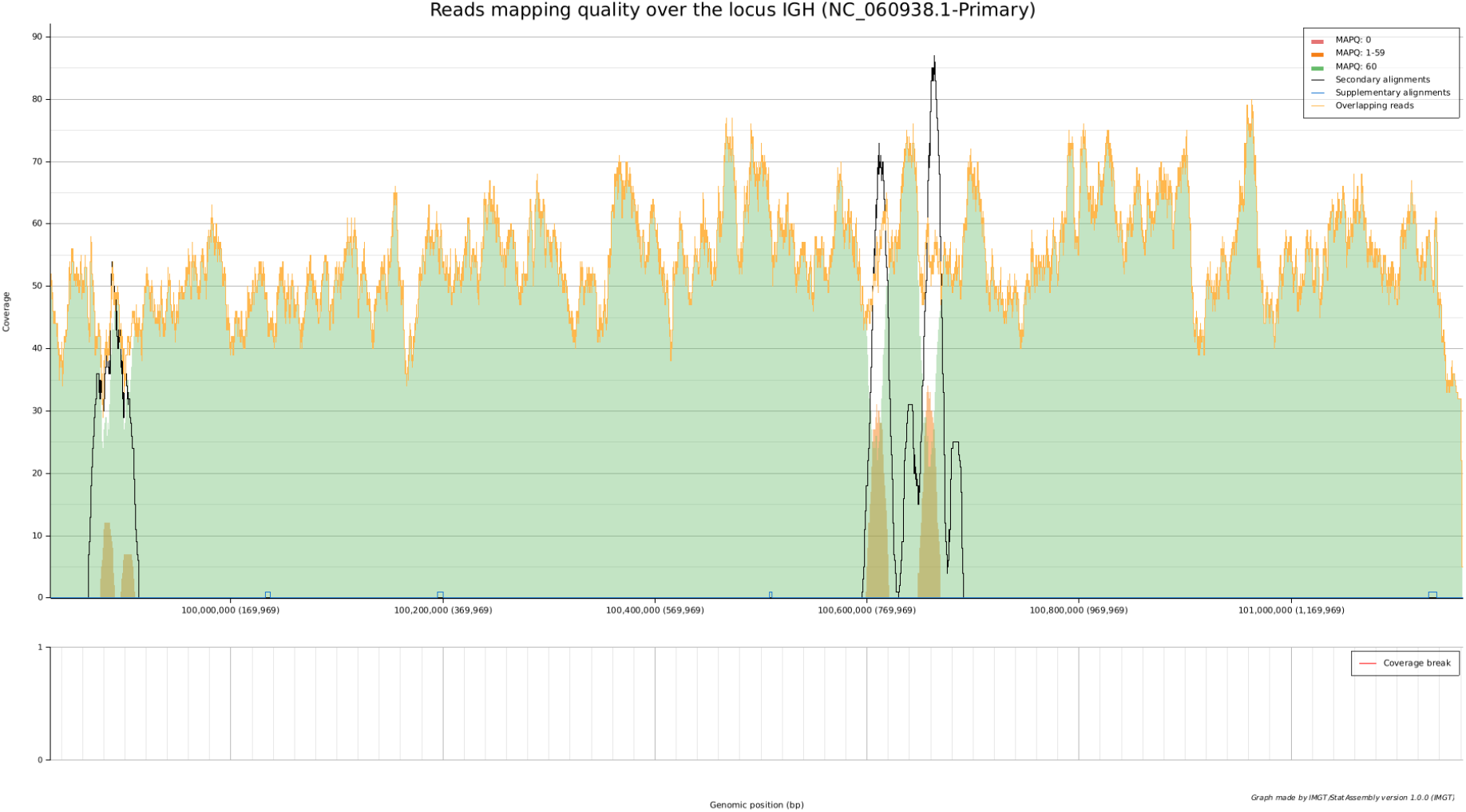
This figure shows the coverage of reads according to their mapping score on the IGH locus for the human T2T-CHM13v2.0 assembly. Upper: The number of primary reads with a mapping quality of 60 (green), between 1 and 59 (yellow), and 0 (red) is counted for each position on the locus. The number of secondary alignments for each position is shown in black, and the number of supplementary/chimeric alignments in blue. Overlapping reads are shown in yellow. Regions where secondary alignments exceed primary ones are the zones around the constant genes (left) and some V genes (right), respectively called CNV7 and CNV3 [13]. Bottom: If the number of primary reads (in green, yellow and red) goes below 3, the ‘break’ region is shown by a red bar. None are observed in this assembly.

With IMGT/StatAssembly, users can also visualize the PHRED score^3^ (upper graph, right axis) and the mismatch ratio (number of primary reads matching the base divided by the number of primary reads aligning at this base) on the top graph (left axis). The mismatch rate for the entire read is displayed in the lower graph (Figure 6). A high-quality assembly should exhibit a high PHRED score (High-Fidelity reads/HiFi) and a lower mismatch rate. In this case, the average mismatch rate does not exceed 0.5%, with an average of less than 10% of mismatches at each position. The average PHRED score is 80.

**Figure 6:**
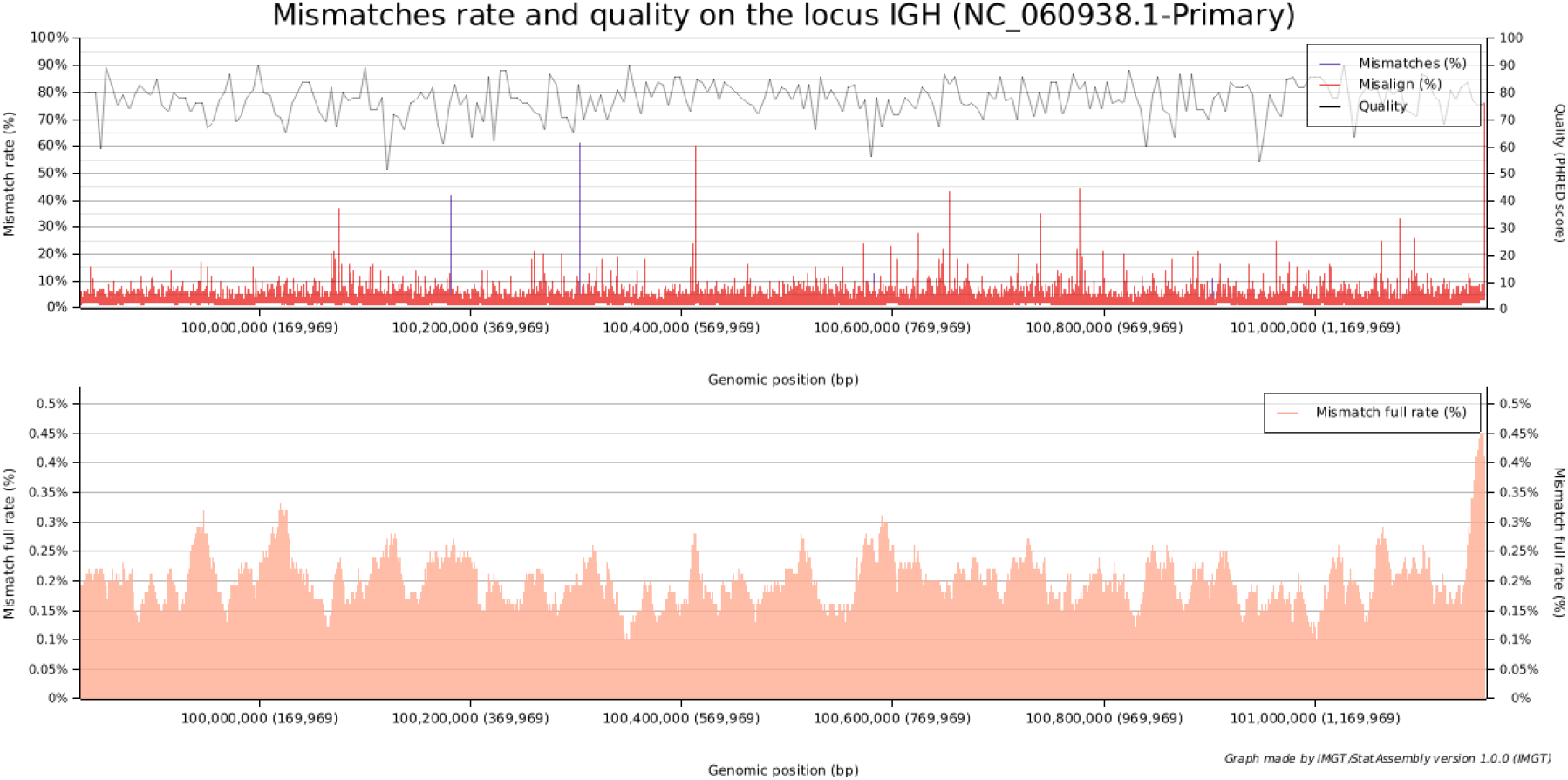
Figure showing the PHRED score and mismatch rate per position (upper) and the mismatch rate over the whole read (lower) on the IGH locus for the human T2T-CHM13v2.0 assembly.

Finally, if gene positions are provided, the number of reads is displayed (marked 1), including matching reads (substitution without indels/marked 3) and identical reads (marked 2) for the precise gene positions, based on the reference sequence. By default, positions with fewer than ten identical reads or an identical-to-total read ratio below 80% are flagged as warning positions and highlighted in yellow. Positions where this ratio drops below 60% are considered suspicious and marked in red. The black curve (marked 4) represents the number of reads that perfectly match (100% match) across this entire region (Figure 7). A confident allele is defined as one with the highest number of reads that perfectly match the reference sequence relative to the total number of reads. The tolerance threshold can also be assessed: both sequence match and sequence identity should fall within 20% of the total number of reads, without triggering any warnings (unless fewer than 10 matching reads are present) or indicating suspicious positions.

**Figure 7:**
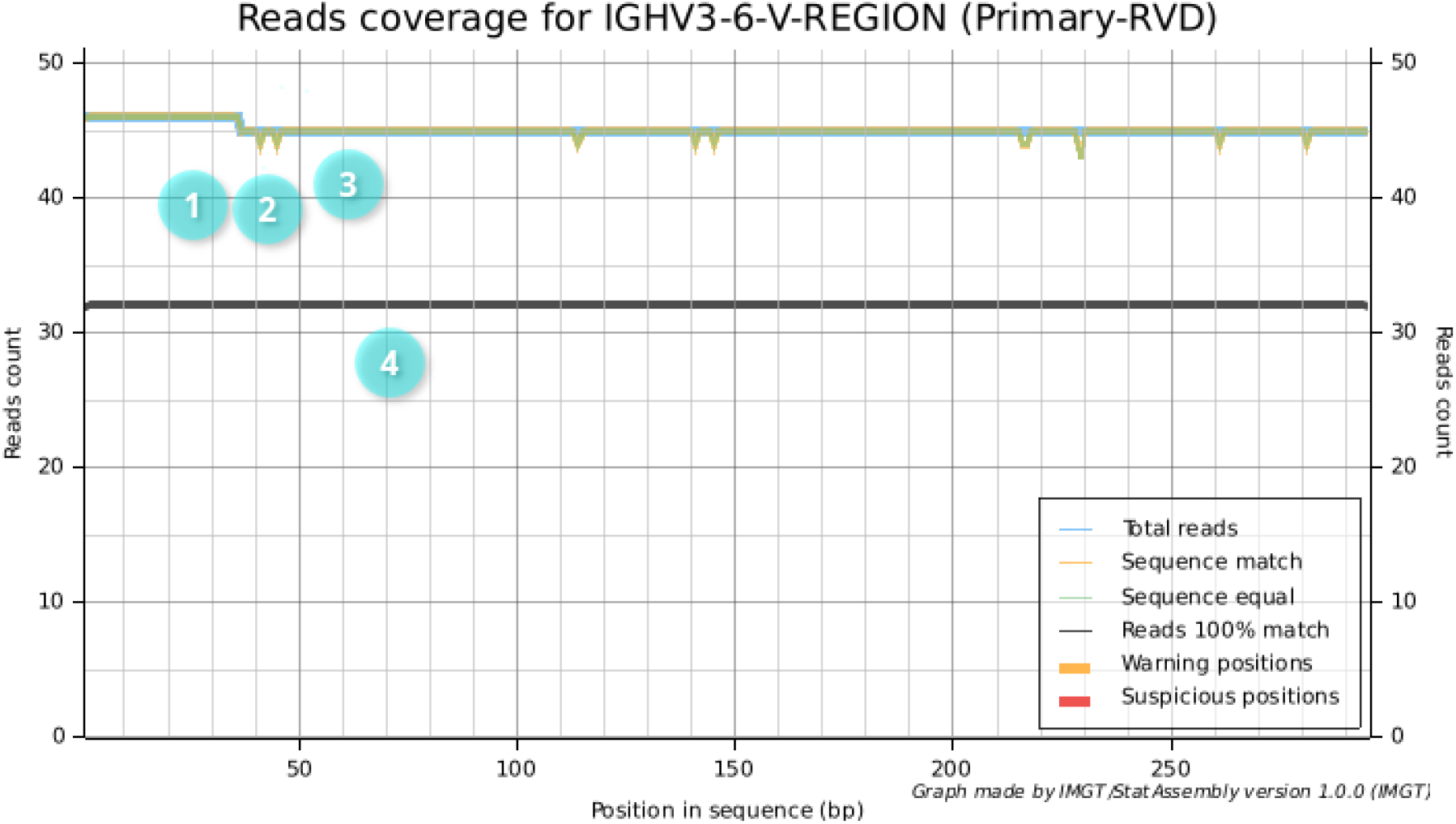
Number of reads aligned over the IGHV3-6 gene V-REGION (NC_060938.1 complement(100326646..100326940)) for the human T2T-CHM13v2.0 assembly. The blue curve (1) represents all reads aligned at each position within the IGHV3-6 V-REGION. The green curve (2) shows reads matching the reference base, while the dark blue curve (3) indicates reads with substitutions at that position. The black curve (4) represents reads that perfectly match the entire IGHV3-6 V-REGION.

Using IMGT/StatAssembly, users can also download these results as CSV files. Regarding genes, a file with multiple columns is generated (see supplementary data *human_IGH_NC_060938*.*1-primary_geneanalysis*).

In conclusion, IMGT/StatAssembly provides an effective means to assess the quality of a genome assembly based on read data. User-defined thresholds can be set as parameters to customize the analysis. The tool also enables evaluation of allele confidence based on read support. By combining IMGT/StatAssembly with the established [IMGT assembly quality rules](https://imgt.org/IMGTScientificChart/Assemblies/IMGTassemblyquality.php), both the overall assembly quality and the reliability of the associated alleles can be systematically assessed. This procedure has been adopted and incorporated in the IMGT standardized biocuration workflow.

#### 2.5.2 Comparaison with CloseRead

Although CloseRead produces assembly-quality results, it does not evaluate the quality of reads for specific genes or alleles. In fact, CloseRead considers a read to be full-length only if there are no mutations across its entire length, excluding indels. However, due to the intrinsic error rate of sequencing reads, this strategy is often impractical. Interestingly, we calculated nine reads that matched the entire IGHA1 C-GENE-UNIT at 100%, whereas CloseRead counted 25. For IGHV3-22, CloseRead reported 34 reads with a perfect match, while we counts 30. For IGHJ2, we found 41 matching reads, whereas CloseRead counted only 27. This discrepancy arises because the mismatch error rate (2, 2 *×* 10^*−*3^) is approximately ten times higher than the mismatch rate (1, 87 *×* 10^*−*4^) in this assembly.

Consequently, for short genes, IMGT/StatAssembly counts are generally higher, whereas for long genes, our counts are typically lower. Therefore, we consider it of paramount importance to recalculate and re-evaluate misalignment and mismatch rates. *We define a read as perfectly mapped if it contains no mismatches or indels within the given gene region*. This re-evaluation was made possible by modifying the library used for BAM file analysis, as the current libraries (hts-lib in Rust and pysam in Python) do not provide this functionality. The resulting library, named extended-htslib, is available under the MIT license. This represents a significant advantage of IMGT/StatAssembly over CloseRead, as it is essential for the accurate validation of new alleles by IMGT rules.

Based on the treatment speed, CloseRead pipeline does not use indexes and can therefore take several hours, depending on the size of the BAM files. In contrast, IMGT/StatAssembly uses indexing and is implemented in a compiled language, allowing it to perform analyses in seconds for assemblies with a lower mismatch rate.

If we consider the multiple mapping, Minimap2[18] may encounter reads that can be mapped to multiple positions. In such cases, the primary alignment is typically assigned a lower mapping quality, as reflected in both tools. However, CloseRead discards the alternative alignments, known as secondary or supplementary alignments. Since these can reveal [potential assembly errors](https://samtools.github.io/hts-specs/SAMv1.pdf), we chose to display and count them. By default, they are shown only in the coverage graph (Figure 5).

Finally, IMGT/StatAssembly generates a dedicated graph for each gene (Figure 7) illustrating the number of matching and non-matching reads, along with the count of perfectly aligned reads. This representation offers a more intuitive and detailed visualization compared to CloseRead.

#### 2.5.3 Perspectives

Our rules evaluate assemblies and alleles in IG/TR loci to ensure consistency, completeness and accuracy of our databases. However, for some assemblies, the raw data may be sporadic or inaccessible. We therefore encourage a collective effort to make assembly sequencing and construction more transparent and standardized, so the scientific community can obtain the highest-quality assemblies possible. Hifiasm and Verkko are commonly used haplotype-resolved assemblers; Hifiasm is based on string graph, whereas Verkko and LJA are based on Bruijn graph.However, no new version of LJA has been released in over three years. Some consortia like Vertebrate Genomes Project [20] aim to achieve this goal and describes procedures for assembly construction. Complete and error-free genomes are particularly important for research and medicine, especially with for IG/TR loci.

IMGT/StatAssembly provides information about assembly quality. However, several additional criteria must be taken into account and are part of IMGT assembly quality rules-such as the number of assemblies with the same gene order, the sequencing technology used, and the assembly method applied. Confidence in the accuracy of alleles and assemblies is higher when high-quality long reads (e.g. HiFi) [21], haplotype-resolved assembly methods [22], and complete genome assemblies such as telomere-to-telomere (e.g. T2T) assemblies [23] are used. Assemblies such as [T2T-CHM13v2.0](https://www.ncbi.nlm.nih.gov/datasets/genome/GCF_009914755.1/), which are derived from the haploid cell line CHM13htert, provide a complete haplotype but do not reflect human genomic diversity [24].

While huge progress has been made in generating complete assemblies[23], challenges remain in producing high-quality assemblies [25]. Moreover, some vertebrate species still have very few assemblies available. A balance must be achieved between assembly availability, sequencing methods, assembly construction, and haplotype resolution in order to accurately capture the full extent of genetic diversity in jawed vertebrates while minimizing false positives.

## 3 AXIS II - IMGT tools for the Analysis of IG and TR repertoires

### 3.1 IMGT/GeneFrequency

[IMGT/GeneFrequency](https://www.ncbi.nlm.nih.gov/datasets/genome/GCF_009914755.1/) is an interactive tool that dynamically evaluates, from IMGT resource [8], the usage of IG and TR variable (V), diversity (D), joining (J) and constant (C) genes expressed in rearranged cDNA sequences and for which the molecule recognized by the antigen receptor may be known. On one hand, this tool allows us to highlight how frequently a gene is used; on the other, for a given specificity, it shows which genes contribute to the synthesis of the molecules that recognize it.

The V, D, J and C genes and alleles that code the IG and TR are managed in IMGT/GENE-DB [5] according to the concepts of Classification of IMGT-ONTOLOGY [3; 26; 27]. IG and TR cDNA that have been submitted to public databases (GenBank, EMBL) are integrated and annotated in the IMGT nucleotide sequence database, IMGT/LIGM-DB [2]. The annotation includes the identification of the involved IG or TR genes and alleles, the identification of the keywords (concepts of identification [28]), the description of the features with IMGT labels (concepts of descriptions [29]) and of their delimitation according to the IMGT unique numbering [30]. Annotation of cDNA rearranged sequences in IMGT/LIGM-DB can be done automatically for productive sequences with a classical organization by IMGT/Automat [31], or semi-manually by biocurators. In the latter case, the specificity is indicated when known.

The web interface of IMGT/GeneFrequency through IMGT-KG [10] query page, is organized in two panels (Figure 8). The right panel is dedicated to the selection of the genes for a given locus and displays the results. The left panel provides a multi-selection parameters, allowing users to select the species, the locus, the specificity, and gene type. The lower part of the left panel lists the available display options, with multiple choices options : bar plot, gene with the known position in the locus only, gene summary table, gene and specificity table, and specificity heatmap. By default, when loading the IMGT/GeneFrequency query page, the bar plot is displayed for the number of cDNA sequences expressing genes of the *Homo sapiens* IGH locus, for the four gene types, regardless of the sequence specificity (Figure 8).

**Figure 8:**
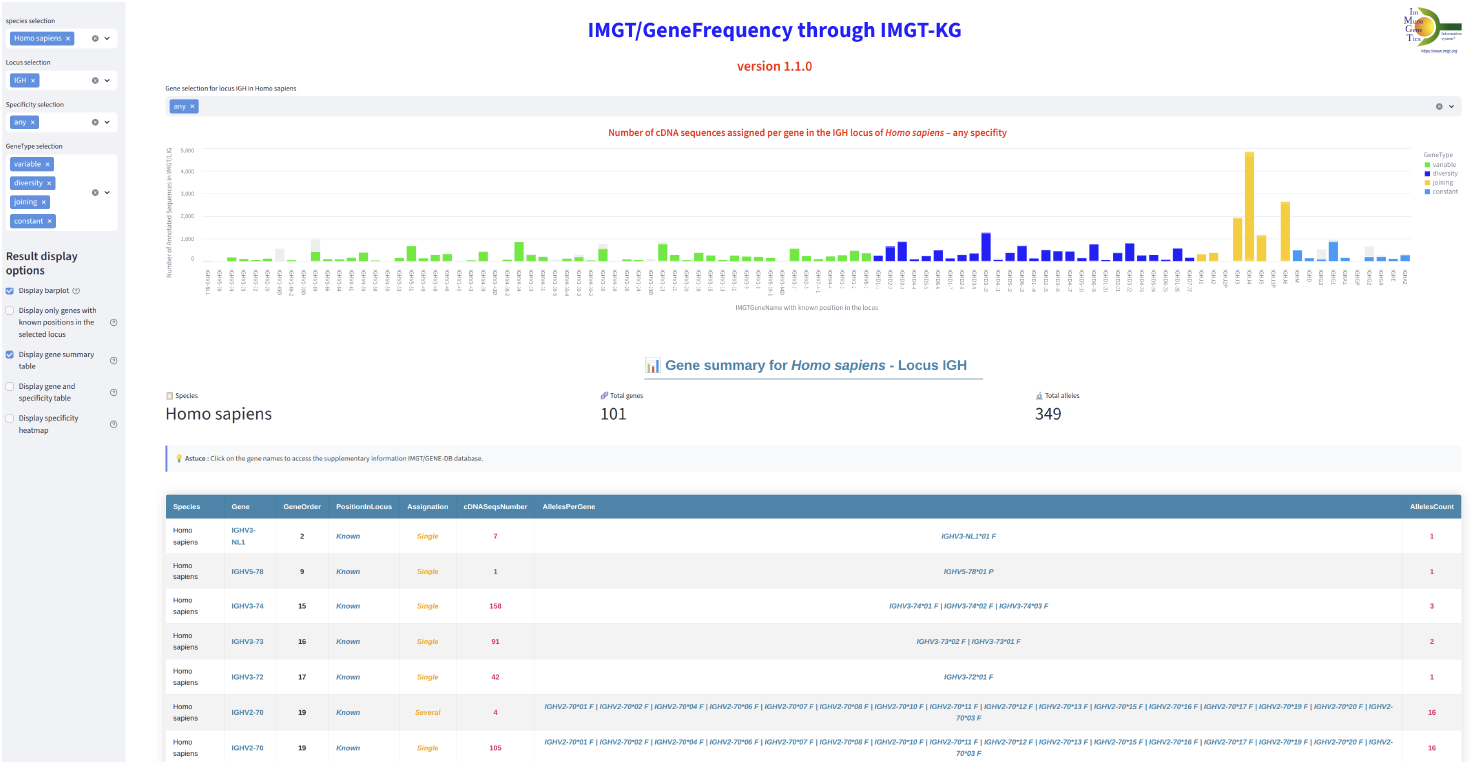
Number of cDNA sequences assigned per gene in the IGH locus of *Homo sapiens* for any specificity

The bar plots are provided per species, locus and specificity. On the x-axis, we have IMGT gene names ordered by their IMGT gene order in the genomic locus. However, non localized genes are listed on the far left part of the bar plot. In order to focus on localized genes only, the user might select the option “Display only genes with known position in the selected locus”. One the y-axis, the number of annotated cDNA sequences in which the genes are expressed is displayed. Only genes expressed in cDNA are shown.

The bars for V, D J and C genes are colored in accordance with the IMGT Color Menu for genes). Note that the bar for a given gene can comprise two parts : a colored part for the number of sequences for the gene which was identified without ambiguity, and a gray part which indicates that the sequence could be assigned to other genes of the same type (for example due to somatic mutations for IG, extensive trimming and or partial sequences). Hovering the mouse on the colored or gray bar displays information about the gene name, gene type, the gene order and the number of annotated sequences.

The results can be displayed in tables per species and IMGT group in which the genes are sorted by their IMGT Gene Order. The **gene summary** table indicates the number of IMGT/LIGM-DB sequences in which it is expressed as well as the names of expressed related alleles and their corresponding number. The table may comprises two lines per gene : one for number of sequences in which it is expressed unambiguously (Single, for single gene), one for sequences where several possibilities exists (several, for several genes). The **gene and specificity** table indicates for each gene pair, the specificity, the number of sequences in which it is identified to be expressed non-ambiguously (Single gene assigned seq), ambiguously (Several genes assigned seq nb (white part), and the total.

Clicking on a gene name in tables displays in a new page the “Table of annotated IMGT/LIGM-DB cDNA rearranged sequences” (Accession number, Allele name, Sequence length, Sequence definitions, Specificity(ies)) provided by IMGT/GENE-DB for that gene. Direct links to IMGT/LIGM-DB entries allow to display the sequence annotations according to the IMGT-ONTOLOGY and IMGT/ScientificChart standards.

Finally, the **Specificity heatmap** evaluates per pair of gene and specificity, and through a color the range of the number corresponding cDNA. Bar plots and heatmaps can be downloaded in SVG or PNG format, where as tables can be exported as CSV files.

IMGT/GeneFrequency results are dynamically computed from IMGT-KG, and will be updated with the evolution of the content of IMGT-KG and the underline databases IMGT/GENE-DB and IMGT/LIGM-DB regarding human and mouse species.

To summarize, IMGT/GeneFrequency based on IMGT-KG, provides an overview of the IG and TR gene expression through the annotated cDNA sequences of IMGT/LIGM-DB, which is enriched with the annotated nucleotides sequences from INSDC (GenBank or ENA). The tool will be available for other species if the number of cDNA sequences integrated in IMGT/LIGM-DB is evaluated as significant to represent the expression of the genes of a given locus. Following the same requirements (sufficient sequence representation), granularity of the results could also include the expression of genes at the allele level. In this current version, it should be noticed that IMGT/LIGM-DB does not include rearranged gDNA or cDNA sequences sets from SRA.

### 3.2 IMGT/V-QUEST: Customize the Reference Directory Set Feature

IMGT/V-QUEST is the IMGT’s flagship bioinformatics tool, specifically designed for the accurate analysis of IG and TR rearranged nucleotide sequences [32]. IMGT/V-QUEST empowers academic researchers and clinicians by providing standardized, high-quality immunogenetic annotations of rearranged IG and TR sequences enabling the study of expressed repertoires in both normal and pathological contexts. The tool aligns and compares the user-submitted sequences with the IMGT reference sequences of V, D and J genes and alleles curated from the IG and TR loci for all supported species and strains.

The IMGT reference directory can be modified or customized based on allele functionality, the inclusion or exclusion of orphon gene sequences and the number of alleles to be considered (all alleles or “01” alleles only) (Figure 9). The restriction of the IMGT reference directory for inbred animal models to sequences from a given strain (Arlee or Swanson for trout IGH and TRB, C57BL/6J strain for mouse IG and TR) has also been progressively introduced.

**Figure 9:**
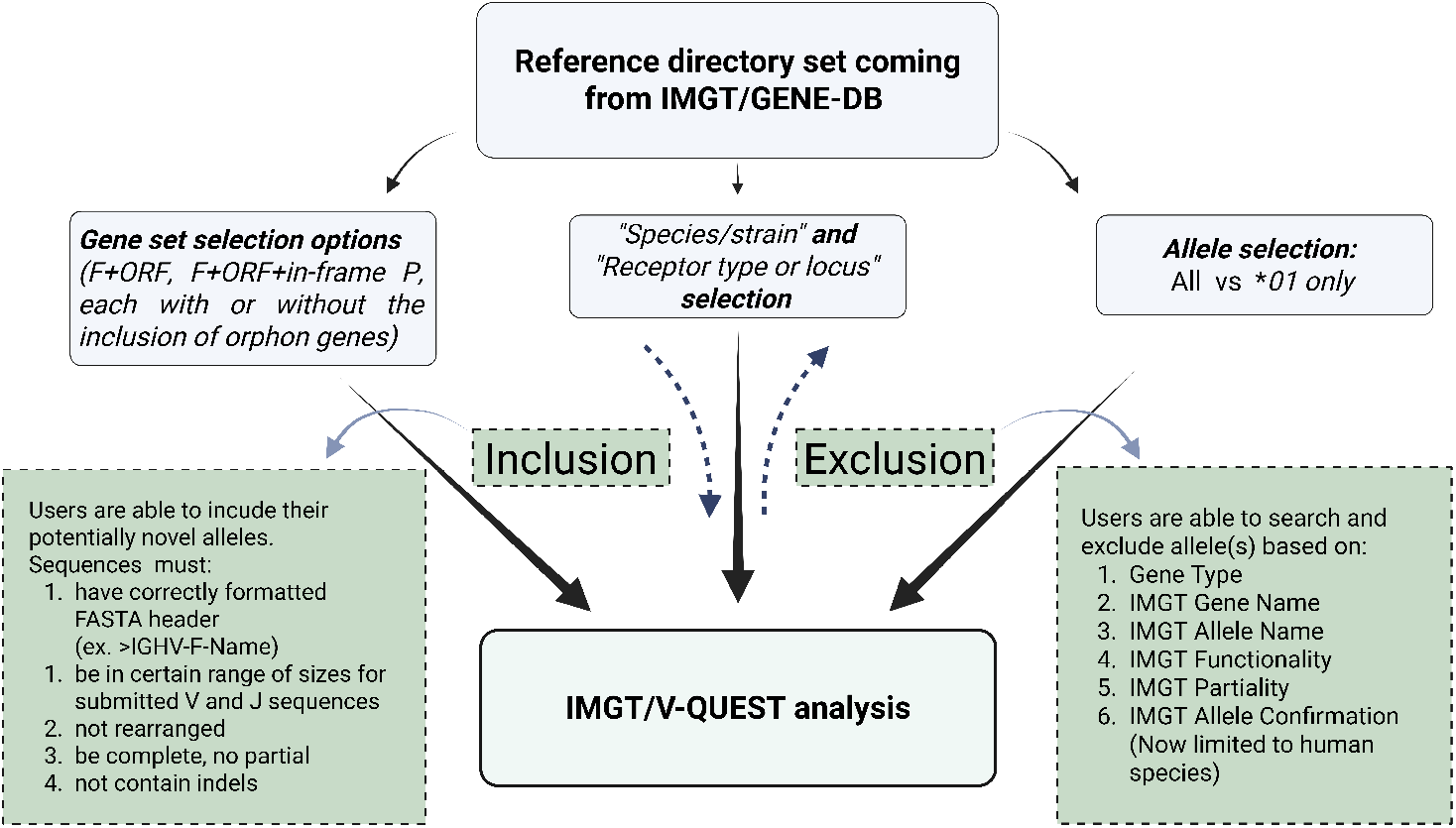
Evolution of reference directory management in IMGT/V-QUEST. The default reference directory set is derived from IMGT/Gene-DB and can be refined through species/strain and receptor type or locus selection, gene set options, and allele selection criteria. Newly introduced features (in dashed green boxes) enable user-driven inclusion and exclusion of specific alleles, enhancing customization and flexibility.

Starting with version 3.7.0, as part of ongoing efforts to enhance the flexibility and user-centric functionality of IMGT/V-QUEST, we have introduced a major new feature that allows greater personalization of the IMGT/V-QUEST reference directory and analysis protocol. This feature, accessible through the “Advanced parameters”, offers users two important customization options of the IMGT/V-QUEST reference directory: (i) selective exclusion of allele reference sequences from standard IMGT reference directory and (ii) inclusion of FASTA sequences representing potentially novel alleles provided by the user (Figure 9 and 10) [33; 8].

**Figure 10:**
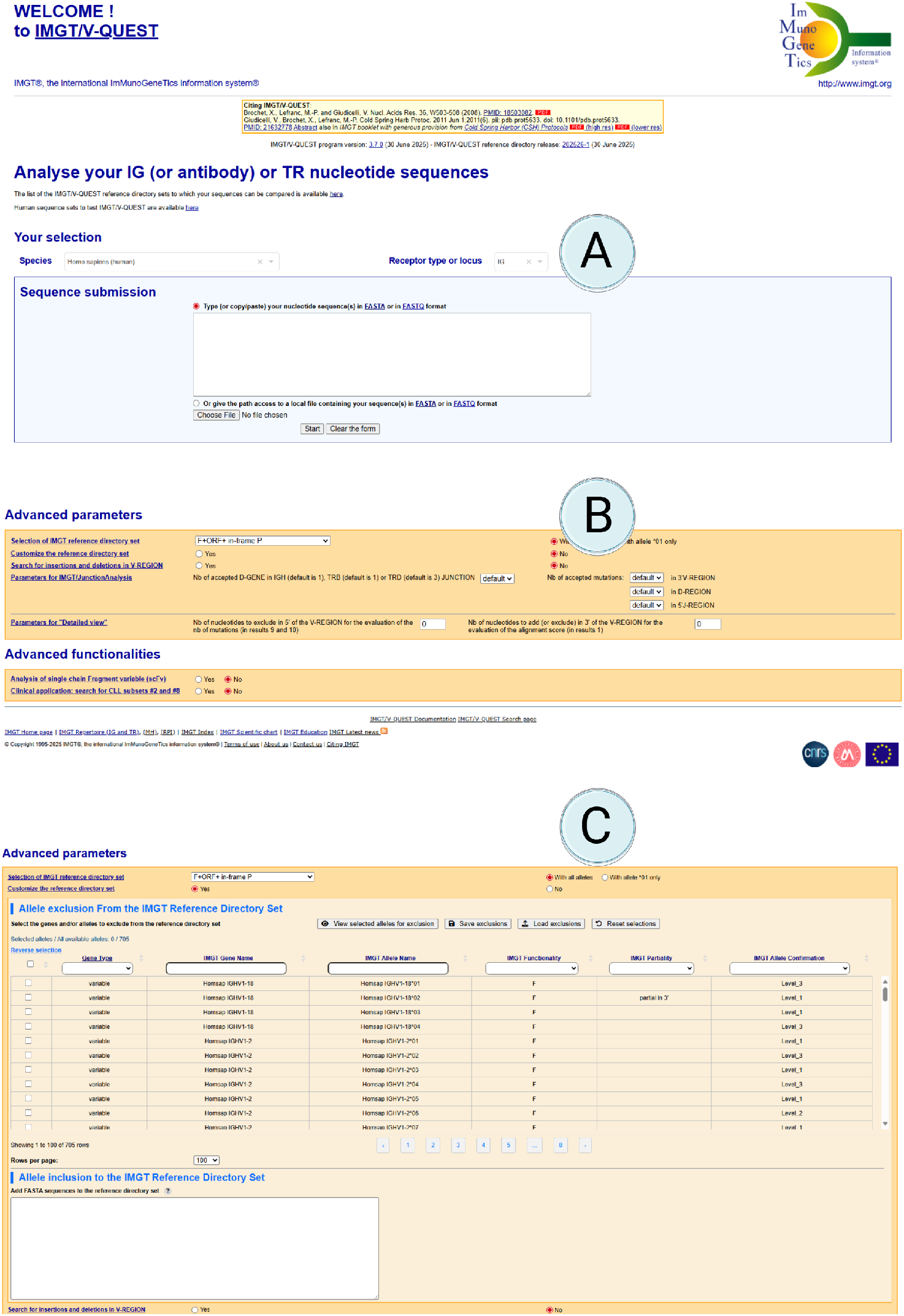
The IMGT/V-QUEST interface illustrating the steps to activate the ‘Customize the reference directory set’ feature: (A) Select the species and receptor type or locus; (B) Enable the customization of the reference sequence directory; (C) Exclude and/or include specific sequences from the IMGT/V-QUEST reference directory set.

This option allows users to control the reference pool by excluding specific sequences from the standard IMGT reference dataset. Exclusion can be based on **Gene Type, IMGT Gene Name, IMGT Allele Name, IMGT Functionality (e.g**., **F, ORF, Pseudogene), IMGT Partiality** for incomplete V, D, or J-REGION, and - for human datasets - **IMGT Allele Confirmation** (evaluating the number of IMGT genomic sequences in which the allele was confirmed).

This option enables scenarios where a smaller or more concentrated reference dataset is required, such as removing pseudogenes or alleles not sufficiently confirmed for a particular study (Figure 10, C). Notably, the user cannot select all the available alleles for a certain chain or gene to exclude, to maintain consistency with IMGT’s reference standards.

In addition, users can now enter and integrate their own custom nucleotide sequences in FASTA format directly into the IMGT/V-QUEST analysis pipeline. This new option significantly enhances the tool’s usability, allowing users to explore the addition of potential new genes and/or alleles to the current IMGT reference database [5; 14].

Researchers working on highly specialized or individualized repertoires will benefit from the ability to analyze custom sequences within the trusted and familiar framework of the IMGT/V-QUEST. To ensure accurate interpretation, all submitted sequences must represent unrearranged germline configurations, fall within the expected length range for V and J gene types, and be free of truncations, partial regions, or indels when aligned with the IMGT unique numbering framework [14].

Furthermore, to ensure system integrity and optimal performance, a limit has been placed on the number of external reference sequences that can be submitted per analysis. This restriction has been introduced to balance between end-user flexibility, computational efficiency, and guarantee a consistent experience across different use cases.

Taken together, these enhancements represent a significant milestone in personalized immunogenetics research, providing users with greater flexibility preserving the robustness of the IMGT/V-QUEST system.

To enhance reproducibility of the IMGT reference directory modification, IMGT/V-QUEST now provides functionality for saving the user-defined reference set configuration, including all the excluded alleles. Users can export their allele exclusion lists, enabling the exact reference customization to be saved, shared, or reloaded for future analyses. In addition, the output report explicitly documents modifications of the reference directory. This ensures full traceability of the analytical context and supports standardized interpretation and comparison of results across studies.

Users are advised to exercise caution when excluding sequences from the reference directory of IMGT/V-QUEST or including user-defined FASTA sequences. Changing the reference dataset either by exclusion or inclusions can significantly affect the results of analysis by IMGT/V-QUEST, with potentially impacting on gene assignment, alignment values, and downstream interpretation. It is the user’s responsibility to ensure that their custom reference set is biologically appropriate and relevant to their specific use case.

To ensure user privacy and data confidentiality, IMGT/V-QUEST does not store or retain any sequences uploaded by users. Submitted user-defined FASTA sequences are retained only for the duration of the session and are automatically removed upon completion of the analysis.

In conclusion, “customize the reference directory set” feature of IMGT/V-QUEST provides the user with greater control and analytical flexibility, enabling customized analyses through the exclusion of specific alleles or the inclusion of user-provided sequences. This capability enhances reproducibility, transparency, and relevance across a wide range of research and clinical applications. The same functionality is planned for future integration in IMGT/HighV-QUEST.

## 4 AXIS III - IMGT tools for Structural Analysis of Immunoproteins

The Axis III provides tools to analyse structural immunoproteins including 2D structures, 3D structures and therapeutic mAbs. In this axis, we introduced IMGT/mAb-KG, a knowledge graph for exploring therapeutic mAbs [12] and IMGT/RobustpMHC, a robust machine learning tool trained for class-I MHC peptide binding prediction [9].

IMGT/mAb-KG is the IMGT-KG specialised in representing and describing therapeutics proteins like mAbs, their therapeutic use and their related clinical indications. IMGT/mAb-KG provides access to over 139,629 triplets describing 1489 mAbs and related proteins, approximately 500 targets and over 500 clinical indications [12]. In addition, IMGT/mAb-KG describes mAbs with their mechanism of action, their construction and the various clinical studies associated. Linked with IMGT-KG, IMGT/mAb-KG provides external links to other domain resources including Thera-SAbDab^4^ [34], PharmGKB^5^ [35], PubMed, and HGNC (HUGO Gene Nomenclature Committee), positioning IMGT/mAb-KG as an essential resource for mAbs engineering. To access to IMGT/mAb-KG, users have two options, one is the use of a SPARQL query interface language and the other is the use of the [exploration interface](https://imgt.org/mAb-KG/). While the first access option may be limited to SPARQL experts, the second offers an intuitive way to explore knowledge about mAbs, including their targets, clinical indications, and modes of action, through graph visualization.

The accurate prediction of peptide-MHC class I binding probabilities is a critical endeavor in immunoinformatics, with broad implications for vaccine development and immunotherapies. For this sake, we have developped IMGT/RobustpMHC that harnesses the potential of unlabeled data in improving the robustness of peptide-MHC binding predictions through a self-supervised learning strategy. The tool is available at https://www.imgt.org/RobustpMHC/ and propose to predict peptide-MHC I and II binding. In each prediction page, the user can either upload his sequences in fasta file or can select the existing HLA sequences as example, then run the predictions. The results consist of a list of HLA and Peptide with prediction values which can be downloaded as a CSV.

## 5 IMGT-KG : a Knowledge Graph for immunogenetics Data

Nowadays, the knowledge graphs (KG) appear as one of the most effective methods for data or knowledge integration and federation. They have gained widespread acceptance in both academic and industry circles [36]. In fact, the complex and interconnected nature of immunogenetics, along with the need to perform sophisticated queries to address complex or integrative biomedical research questions, has rapidly accelerated the adoption of knowledge graph technologies[37].

To unify and connect various IMGT resources, we developed IMGT-KG, the first Findable, Accessible, Interoperable, and Reusable (FAIR) Knowledge Graph (KG) in immunogenetics, providing access to structured and enriched immunogenetic data [10]. IMGT-KG bridges the gap between nucleotide and protein sequences of IMGT databases. For that, IMGT-KG acquires data from IMGT databases, then represents, describes and structures immunogenetics entities and their interrelationships in a KG using semantic web standards and technologies. In addition, IMGT-KG is connected to external resources in the same domain, such as the Relation Ontology (RO) [38], Feature Annotation Location Description Ontology (FALDO) [39], NCIt [11], Sequence Ontology (SO) [40], and other IMGT-KG opens the way for effective queries and integrative immuno-omics analyses over more than 100 million triplets. Users can access these two IMGT-KG using the SPARQL language through https://imgt.org/imgt-kg/.

## 6 Conclusion

To decipher the complex mechanisms behind the adaptive immune responses and the associated diversity of the involved receptors, we assisted in a massive generation of immunogenetics data at different levels (genomics, transcriptomics, proteomics, …) thus plunging the immunogenetics domain into the era of Big Data [41; 1; 42]. To manage, analyse, and interpret this rich immunogenetics data, IMGT has set up different databases, tools and resources from the nucleotide sequences to protein sequences for analysing, visualizing, and interpreting these data through three axes [8]. Axis I aims to decipher the adaptive immune response by analysing the IG and TR loci across jawed vertebrates. Axis II investigates IG and TR gene diversity and the expression patterns essential to understanding the adaptive immune response in normal and pathological situations. Axis III analyses the 2D and 3D structures of the adaptive immune proteins to enhance our understanding of the molecular mechanisms that drive immune responses.

In response to the growing and evolving needs of the immunogenetics research community, IMGT has enhanced its existing tools and resources and developed new ones, either starting from scratch or by collaborating with the scientific community to build upon existing tools. In fact, in Axis I, new user-friendly, dynamic gene tables have replaced time-consuming and error-prone gene tables and CDR length tables. New tools for analysing assemblies, such as IMGT/AssemblyComparison and IMGT/MGV, have been created. Axis I also implements a set of rules and a new tool: IMGT/StatAssembly, which validates the quality of loci and alleles in light of the large-scale generation of immunogenetics data.

In the Axis II, we introduced a new generation of IMGT/GeneFrequency, with a modern interface and new features and functionalities. The new IMGT/GeneFrequency is based on the IMGT-KG [10] and introduces different visualisation (barplot and heatmap) and tables (gene summary, gene and specificity). We also enhanced IMGT/V-QUEST by introducing the possibility to customise the reference directory set using the gene type, the gene and the allele name, the allele functionality, and the allele partiality. This new feature also allows the addition of the user sequence in the IMGT/V-QUEST analysis pipeline, thus enhancing the result for new genes and alleles.

In the Axis III, we introduced IMGT/RobustpMHC [9], a new robust machine learning tool for peptide-MHC prediction. Based on self-supervising learning strategy, IMGT/RobustpMHC aims to boost the immunotherapies and vaccines development.

To enhance IMGT data accessibility and integrative immunogenetics analyses, IMGT has developed IMGT-KG [10], the first FAIR KG in the immunogenetics field bridging the gap between nucleotide and protein sequences of IMGT database and connected also IMGT resources to external knowledge in the biomedical domain. In addition, to IMGT-KG, we introduced lastly a specialised KG dedicated to the therapeutic monoclonal antibodies : IMGT/mAb-KG. Built on top of semantic web technologies, both IMGT-KG and IMGT/mAb-KG provide facilities to access IMGT immunogenetics data through an efficient and powerful query language (SPARQL) and an intuitive visual query interface in the case of IMGT/mAb-KG. Using the semantic web principles, IMGT-KG and IMGT/mAb-KG play an essential role in the standardisation and the opening of the immunogenetics data on the web thus placing IMGT and its related axes in the world of Linked Open Data [43].

## Supporting information

human_IGH_NC_060938.1-primary_geneanalysis

## 7 DATA AVAILABILITY

IMGT® is freely available online for academics and non-profit use at https://www.imgt.org/. All the resources referred to in this article are accessible from IMGT® webpages. IMGT/StatAssembly source code and documentation are available on Gitlab (https://src.koda.cnrs.fr/imgt-igh/statassembly).

## 8 ACKNOWLEDGEMENTS

We would like to thank Anthony Boureux for insightful discussions on k-mers as well as Christophe Klopp, members of Beijing Institute of Genomics (BIG) and Corey Watson for their detailed input on assembly reconstruction and haplotypes. We are also grateful to Yixin Zhu and Anton Bankevich for their helpful discussions on the CloseRead software. We thank Hervé Seitz’s team at Montpellier MGX GenomiX for valuable exchanges regarding sequencing and genome assembly reconstruction. Finally We thank Dr. Anna Tran at MéLiS (UCBL)/Hospices Civils de Lyon for her initial work and Joel Richardson from MGI for sharing the MGV code with us.

We thank all members of the IMGT® team for their expertise and constant motivation. IMGT® is member of the French Infrastructure Institut Français de Bioinformatique (IFB) as well as member of BioCampus, MAbImprove and IBiSA.

## Footnotes

1 Secondary alignments are alignments of reads of the lowest quality, due to repeats in the assembly, as stated by minimap2 article.

2 Supplementary alignments are additional alignments of a read that cannot be represented as a single linear alignment, typically due to chimeric or split reads, as described in the Minimap2 [18] article.

3 PHRED quality score is used as an indicator of base quality in DNA sequencing

4 Therapeutic Structural Antibody Database

5 comprehensive resource curating knowledge on the impact of genetic variation on drug response

